# ROUA Database: 300 Human Genomes from the border of Ukraine and Romania

**DOI:** 10.1101/2024.02.27.582437

**Authors:** Khrystyna Shchubelka, Walter W. Wolfsberger, Olga T. Oleksyk, Yaroslava Hasynets, Silviya Patskun, Mykhailo Vakerych, Roman Kish, Violeta Mirutenko, Vladislav Mirutenko, Coralia Adina Cotoraci, Calin Pop, Olimpia Neagu, Cornel Baltă, Hildegard Herman, Paula Mare, Simona Dumitra, Horatiu Papiu, Anca Hermenean, Taras K. Oleksyk

## Abstract

We present a multi-layered data source (ROUA Database) providing the results of Whole Genome Sequencing of two human populations in the Carpathian Mountains region, specifically Ukraine’s Transcarpathia and Romania’s Satu Mare and Baia Mare provinces, areas previously underexplored in population genomics. The database contains the raw and annotated files of the whole genome sequences from 300 individuals from these regions, including annotations of common and unique genetic variants following a sampling protocol designed to capture the genetic diversity of Ukrainians and Romanians, including minority groups like Wallachians and Roma. The data is hosted on a dedicated web resource. We provide information on how to access to results of primary and secondary analysis of the data, including comparative analysis with previously published populations from Ukraine, and populations from International Genome Sample Resource and Human Genome Diversity Project. The free research access to this database is contributing to growing understanding of human genetic diversity in Central Europe. This effort emphasizes the potential for reuse of the generated data, advocating for open access to support future research in genomics, bioinformatics, and personalized medicine.

## Data description

The database contains sequences of whole genome from 300 individuals from the two modern human populations living in the Carpathian Mountains region at the international border between Ukraine and Romania, specifically in the Ukraine’s Transcarpathia and Romania’s Satu Mare and Baia Mare provinces, areas previously underrepresented in population genomics databases. The database is located inside a dedicated web portal that contains other materials related to the “*Partnership for Genomic Research in Ukraine and Romania*” project performed by the Ukrainian and Romanian partners and supported by the *Joint Operational Programme Romania-Ukraine, through the European Neighbourhood Instrument* (ENI) (https://genomes.uzhnu.edu.ua/). The portal also provides detailed descriptions of the project, types of data available, and a form to request the access to the data at various analytical stages.

The sampling process for the study was authorized by the Institutional Review Board of Uzhhorod National University, with medical professionals across specific regions supervising blood sample collection in hospitals. Healthy volunteers were recruited through advertisements for interviews at outpatient facilities, where they provided informed consent for their data’s public availability and completed a questionnaire on their background and health history. All personal identifiers were anonymized post-collection, with samples labeled with unique codes. Blood samples were drawn into EDTA tubes, sent to a certified lab for DNA extraction, and any excess stored at Uzhhorod National University’s biobank. In total, 300 participants contributed samples, which were sequenced using the DNBSEQ-G50 platform. The sequencing data were processed using a high-performance pipeline, with reads aligned to the GRCh38 human genome and prepared for variant calling, resulting in a database encompassing Whole Genome Sequencing and analytical results. The resulting database combines the results of Whole Genome Sequencing (WGS) results and primary and secondary analysis results. The raw sequencing reads for every sample in the populations are stored in compressed FASTQ format. Summary sequencing statistics are presented in **Table 1**.

**Table 1.** Summary sequencing statistics and the properties of the raw read files from 300 individuals from the two modern human populations living in the Carpathian Mountains region at the international border between Ukraine and Romania: Ukraine’s Transcarpathia and Romania’s Satu Mare and Baia Mare provinces.

| Population | N. of samples | Read length, bp (average) | Total Reads (average) | Total Bases (average) | Read Q20 (average %) |
| --- | --- | --- | --- | --- | --- |
| *Ukrainians in Transcarpathia* | 150 | 150 | 666,990,234 | 100,046,395,519 | 97.75 |
| *Romanians in Satu Mare and Baia Mare provinces* | 150 | 150 | 674,134,737 | 101,120,210,556 | 96.28 |

Comparative datasets include the results analysis of genome diversity and population contrasts with genetic variation in various Eurasian populations samples from International Genome Sample Resource and Human Genome Diversity Project (Bergström et al., 2020; Fairley et al., 2020). These datasets contain the tabular and graphical results of Principal Component Analysis, Admixture analysis, and Pairwise Fisher Exact Test (FET) analysis for allele frequencies for all the populations included. In addition, we provide access to processed and filtered Variant Call Format of potentially clinically causative alleles, with integrated annotations from ClinVar database.(Landrum et al., 2018). The analysis data includes functionally annotated Variant Call Format (VCF) files, that were processed using an adapted GATK Best Practices pipeline. The summary statistics of the VCF files are provided in **Table 2**.

**Table 2.** Annotation Summaries the 300 genomes dataset. The dataset includes genomes from 300 individuals from the two modern human populations living in the Carpathian Mountains region at the international border between Ukraine and Romania: 150 samples from Ukraine’s Transcarpathia (UA150) and 150 samples from Romania’s Satu Mare and Baia Mare provinces (RO150)

| Number of effects by region |  |  |  |  |
| --- | --- | --- | --- | --- |
| Type (alphabetical order) | RO150 |  | UA150 |  |
|  | Count | Percent | Count | Percent |
| **Downstream regions** | 7,576,139 | 5.55% | 8,694,629 | 9.51% |
| **Exons** | 1,796,755 | 1.32% | 1,403,258 | 1.54% |
| **Genes** | 943 | 0.00% | 2,352 | 0.00% |
| **Intergenic Regions** | 11,380,842 | 8.34% | 10,108,582 | 11.06% |
| **Introns** | 105,645,185 | 77.37% | 61,663,007 | 67.47% |
| **Splice site acceptors** | 7,386 | 0.01% | 5,326 | 0.01% |
| **Splice site donors** | 6,336 | 0.01% | 5,338 | 0.01% |
| **Splice site regions** | 169,779 | 0.12% | 107,995 | 0.12% |
| **Transcripts** | 334,150 | 0.25% | 4,810 | 0.01% |
| **Upstream regions** | 7,495,661 | 5.49% | 8,744,166 | 9.57% |
| **3' UTR regions** | 1,662,611 | 1.22% | 508,075 | 0.56% |
| **5' UTR regions** | 461,771 | 0.34% | 142,714 | 0.16% |

| **Functional Annotation**<br>(# effects by impact severity) |  |  |  |  |
| --- | --- | --- | --- | --- |
| **Type** | **RO150** |  | **UA150** |  |
|  | **Count** | **Percent** | **Count** | **Percent** |
| **High** | 41,966 | 0.03% | 23,203 | 0.03% |
| **Low** | 663,955 | 0.49% | 300,938 | 0.33% |
| **Moderate** | 575,400 | 0.42% | 224,767 | 0.25% |
| **Modifier** | 135,256,237 | 99.06% | 90,841,344 | 99.40% |

| **Number variants by type** |  |  |
| --- | --- | --- |
| **Type** | **RO150** | **UA150** |
|  | **Total** | **Total** |
| **SNP** | 17,884,931 | 18,704,768 |
| **Ins** | 2,607,813 | 2,938,616 |
| **Del** | 2,573,906 | 3,429,057 |
| **Total** | **23,066,650** | **25,072,441** |

## Context

The Carpathian Mountains region in Eastern Europe including Ukraine’s Transcarpathia and Romania’s Satu Mare and Baia Mare provinces, has been under-researched in terms of population genomics despite its rich history and ethnic diversity (Oleksyk et al., 2022). The region’s socio-economic reliance on agriculture and lower levels of industrialization compared to other areas in Ukraine and Romanian provides potential for important genomic discoveries. Previous research focused on mitochondrial haplogroups but lacked comprehensive genome-wide analysis. This study presents a cross-border population genomic analysis, sequencing 300 individuals from Transcarpathia, Satu Mare and Baia Mare provinces. Uncovering unique genetic variants in these culturally and geographically significant areas will contribute to the understanding of human genetic diversity in Central Europe and expand on previously published 97 WGS samples produced in Ukraine (Oleksyk et al., 2021). This research, funded by the European Union under the Joint Operational Program Romania - Ukraine 2014-2020 (Grant ENI CBC-2SOFT/1.2/48), was conducted in collaboration between researchers in the border regions between Romania and Ukraine.

## Methods

The sampling procedure received approval from the Institutional Review Board (IRB) of Uzhhorod National University (Protocol #1 dated 09/18/2018). Sampling was performed with aim to capture the genetic diversity of Ukrainians across all districts of Transcarpathia (Zakarpatska Oblast), and Romainas across Satu Mare and Baia Mare provinces. Additionally, genetic samples from minority groups, specifically Wallachians and Roma, were included.

Medical professionals Transcarpathia and Romania’s Satu Mare and Baia Mare provinces were enlisted to supervise the collection of blood samples in hospital settings. Healthy individuals, not currently hospitalized, were recruited via advertisements and scheduled for interviews at outpatient facilities. During these sessions, participants were briefed on the study’s objectives and the sampling process, providing informed consent for their genotypic and phenotypic information to be made accessible to the public. Participants also filled out a questionnaire detailing their self-reported ancestral background, birthplaces of grandparents (when known), gender, and certain phenotypic traits, including a brief health history. Documentation of consents and interviews are securely archived at the Biology Department of Uzhhorod National University. Following the interview and collection process, personal identifiers were removed from the sample containers, which were then labeled with a unique alphanumeric code and barcode, ensuring anonymity in all further analyses and publications.

After completing the interview, a certified healthcare professional drew a whole blood sample into two 5 ml EDTA tubes, each marked with a barcode, and sent to a certified biomedical lab in, on dry ice for immediate DNA extraction upon receipt. Any remaining blood and DNA samples post-genetic analysis are preserved at the biobank of the Biology Department, Uzhhorod National University, Ukraine and at “Vasile Goldi□” Western University of Arad, Romania.

For all 300 samples, DNA was extracted at Uzhhorod National University’s Molecular Genetics laboratory, using the Monarch DNA purification kit (New England Biolabs, Inc., Rowley, MA, USA) to extract genomic DNA from the original frozen whole blood samples. Approximately 1μg of genomic DNA was fragmented using *Covaris* (Woburn, Massachusetts) and subsequently prepared for DBNSEQ-G50 sequencing at the BGI-Copenhagen in Denmark).

All the individuals in this study were sequenced with DNBSEQ-G50. The sequencing data reads produced using the platform for 300 samples were aligned to the GRCh38 human reference genome using BWA-MEM (Version: 0.7.16a-r1181). Variant Calling was according to the GATK Best Practices Guidelines (Depristo et al., 2011), using a pipeline adapted by us from the Snakemake workflow catalog, (Köster et al., 2021) and hosted on GitHub(https://github.com/valerpok/dna-seq-gatk-variant-calling). Variant calling was performed in two separate batches (150 Ukrainians and 150 Romanians) and merged for subsequent analysis.

Sequence variant files were annotated using SNPEff (SNPE*ff*, RRID:SCR_005191)(Cingolani et al., 2012) software using GRCh38 reference annotation databases. We used *CliVar* (Landrum et al., 2016) and GWAS catalog (Sollis et al., 2023) databases for annotation of the medically related and functional variants using the *snpSift* tool.

The database contains information on genetic diversity and admixture. To perform Principal Component Analysis (PCA), we merged the WGS of our study with European samples from the 1,000 Genomes Project and Human Genome Diversity Project (HGDP)(Fairley et al., 2020). The analysis was conducted using *Eigensoft* (Price et al., 2006). Post-genotyping, rate filtering, and pruning for linkage disequilibrium, 677 samples with 208,945 variants remained. PCA was visualized using Python with pandas, matplotlib, and seaborn libraries, excluding two outlier samples. To perform model-based population structure analysis using the same dataset, we used ADMIXTURE (Alexander et al., 2009) software. We determined the optimal K parameter as 3 through 10-fold cross-validation, which concorded with our previous findings from WGS project in Ukraine. Population structure plots were created using Python, incorporating samples from the IGSR and HGDP databases.

## Reuse Potential and Data Availability

Our database provides free genome data to researchers in Central Europe and is filling and important blank spot in our understanding of European genomic diversity and population history. This database follows open access philosophy, with data released by request for research. The data is hosted on a web resource at Uzhhorod National University (https://genomes.uzhnu.edu.ua/), providing a source for future genomic, bioinformatic, and personalized medicine research.

## Declarations

## Author Contributions

KS organized and supervised collection of the data and wrote the first draft. W.W.W. and K.S. prepared and analyzed the data. All other authors contributed ideas to the development of the pipeline and reviewed the the manuscript. T.K.O. contributed to the original ideas, writing, and final editing of the manuscript.

## Dara Availability

The data with the complete instructions for ROUA Database use, the supplementary materials including the collection protocols, the informed consents, and the decisions of the Institutional Review Board (IRB) of Uzhhorod National University (Protocol #1 dated 09/18/2018 are available through the web portal (https://genomes.uzhnu.edu.ua/),

## List of Abbreviations

WGS: Whole Genome Sequencing;
UA 150: Ukrainians in Transcarpathia;
RO150: Romanians in Satu Mare and Baia Mare provinces;
PGP: PopGenPlayground;
VCF: Variant Call Format.

## Competing Interests

The authors declare that they have no competing interests.

## Funding

Funding for the project was provided by 2SOFT/1.2/48 project “*Partnership for Genomic Research in Ukraine and Romania*” by the *Joint Operational Programme Romania-Ukraine*, through the *European Neighbourhood Instrument* (ENI).

## Acknowledgements

The ROUA database is part of the developing infrastructure for bioinformatics in Ukraine. We thank all the participants of the *BioinformaticsForUkraine*.*com* and the *Genome Diversity in Ukraine Consortium* who worked with us on developing tools for this project.

## Notes

https://genomes.uzhnu.edu.ua/

### Competing Interest Statement

The authors have declared no competing interest.

